# Dynamic interplay between RNA *N*^6^-methyladenosine modification and porcine reproductive and respiratory syndrome virus infection

**DOI:** 10.1101/2024.12.03.626656

**Authors:** Zi-Han Wang, Jing Li, Sai-Ya Ma, Meng-Xuan Liu, Yu-Fei Zhan, Feng Jin, Bing-Xin Liu, Wenjing Wang, Mei He, Yu-Chuan Yang, Yandong Tang, Peng Wang, Wuchao Zhang, Jie Tong

## Abstract

*N*^*6*^-methyladenosine (m^6^A) has attracted significant attention for its role in various biological processes, including RNA stability, translation, and the immune response. Understanding the role of m^6^A in viral infections is crucial to deep the complex interaction between virus and host cells. Porcine Reproductive and Respiratory Syndrome Virus (PRRSV) is a significant pathogen affecting swine health worldwide. Here, we firstly identified the m^6^A peaks in the PRRSV genome by m^6^A RNA immunoprecipitation sequencing (m^6^A-seq). Seven m^6^A-enriched regions within the PRRSV genome were detected, with one located in the N protein-coding region and the others distributed across nonstructural protein-coding regions. Notably, the Nsp2-coding region contained the highest m^6^A peak, spanning approximately 178 nucleotides. Functional analyses demonstrated a positive correlation between m^6^A modification levels and PRRSV replication in porcine alveolar macrophages (PAMs), as modulating the expression of m^6^A methyltransferases and demethylases significantly influenced viral replication. Moreover, treatment with the universal methylation inhibitor 3-deazaadenosine (3-DAA) effectively suppressed PRRSV replication, suggesting its potential as a novel anti-PRRSV therapeutic. To further elucidate the role of m^6^A in PRRSV infection, we analyzed the m^6^A landscape in PAMs infected with pandemic and highly pathogenic PRRSV strains. Among the 4,677 transcripts exhibiting altered m^6^A modification levels, the MAPK14 gene as well as other genes in p38/MAPK signaling pathway potentially emerged as the preliminary targets of m^6^A-mediated epigenetic regulation during PRRSV infection. These findings provide new insights into the epigenetic mechanisms underlying PRRSV infection and may facilitate the development of targeted anti-PRRSV strategies and therapeutics.

**IMPORTANCE:** The involvement of m^6^A in viral genome has broader implications for our understanding of virus evolution and pathogenicity. The ability of viruses to adapt to their hosts and exploit host cellular mechanisms is a critical factor in their success as pathogens. As m^6^A is a modification that can be manipulated by both host and virus, studying its role in viral infections may reveal new insights into viral evolution. Understanding how different viruses utilize m^6^A could inform strategies for developing antiviral therapies that target these interactions. Such therapies could disrupt the ability of viruses to exploit m^6^A modifications, potentially reducing their virulence and enhancing host defenses.

## OBSERVATION

The role of RNA modifications as critical regulators of gene expression has been explored in the context of viral infections during the past several years [1-3]. Among these modifications, *N*^6^-methyladenosine (m^6^A) has attracted significant attention for its role in various biological processes, including RNA stability, translation, and the immune response [4,5]. m^6^A is the most prevalent internal modification in eukaryotic messenger RNA (mRNA), and its dynamic nature allows mRNA for rapid alterations in response to intra- and extracellular insults [6-8]. The significance of m^6^A extends beyond cellular RNA to include viral RNA, as many viruses exploit this modification to enhance their replication and evade host immune defenses [9].

Porcine Reproductive and Respiratory Syndrome Virus (PRRSV) is a significant pathogen affecting swine health worldwide, leading to considerable economic losses in the pig industry [10]. Understanding the mechanisms of PRRSV infection is essential for developing effective control strategies. The emergence of PRRSV was first documented in the United States and Europe in the late 1980s [11]. The situation intensified in 2006 with the epidemic of a highly pathogenic variant of PRRSV (HP-PRRSV) in China, which led to more severe outbreaks characterized by higher morbidity and mortality rates [12-14]. PRRSV NADC30-like strains first emerged in China in 2013 [15]. These strains share genetic similarity with the NADC30 strain isolated in the United States, which load a 131-amino-acid deletion within the hypervariable region of the nsp2 protein [16].

In our study, we utilize the HP-PRRSV HuN4 strain and pandemic NADC-30 like strain HeN-L1. We first employed an ELISA-based approach to assess the presence of m^6^A modifications in PRRSV genomic RNA. Porcine alveolar macrophages (PAMs) were infected with either the PRRSV HuN4 strain or a PRRSV NADC30-like strain, and after 48 hours, the cells were subjected to three freeze-thaw cycles. The supernatants were collected by centrifugation, followed by ultrafiltration to concentrate the viral particles, from which viral RNA was extracted and fragmented into 80-200 nt length. The presence of m^6^A was then measured as a percentage of all adenosines in the viral RNA sample using ELISA. As shown in Figure 1A, the viral RNA had an m^6^A percent of approximately 0.6-0.8%, which double times higher than the 0.3-0.4% observed in total cellular RNA. To further mapping the m^6^A-modified regions within the viral genome, we conducted MeRIP-seq method. PAMs cells were infected with the PRRSV NADC30-like strain, and at 48 hours post-infection, cells were subjected to three freeze-thaw cycles, followed by centrifugation and ultrafiltration to concentrate viral particles. The extracted viral RNA was fragmented and immunoprecipitated with an m^6^A-specific antibody after UV crosslinking. Reverse-transcription and the subsequent deep sequencing of the RNA-antibody complexes were applied to identified the m^6^A modified region in PRRSV genomic RNA. The stringent peak calling method (FDR<0.01) were applied to determine the statistically significant peaks (*p* values <1E-10). Moreover, correlation test between three biological replicates of viral RNA showed a high correlation (0.996), which confirm the replicability of the results. It has revealed seven m^6^A-enriched regions in the viral genome, with one located in the N protein-coding region and the others distributed across nonstructural protein-coding regions. Notably, the Nsp2-coding region contained the highest m^6^A peak, of which the length is approximately 178 nt.

**Figure 1.**
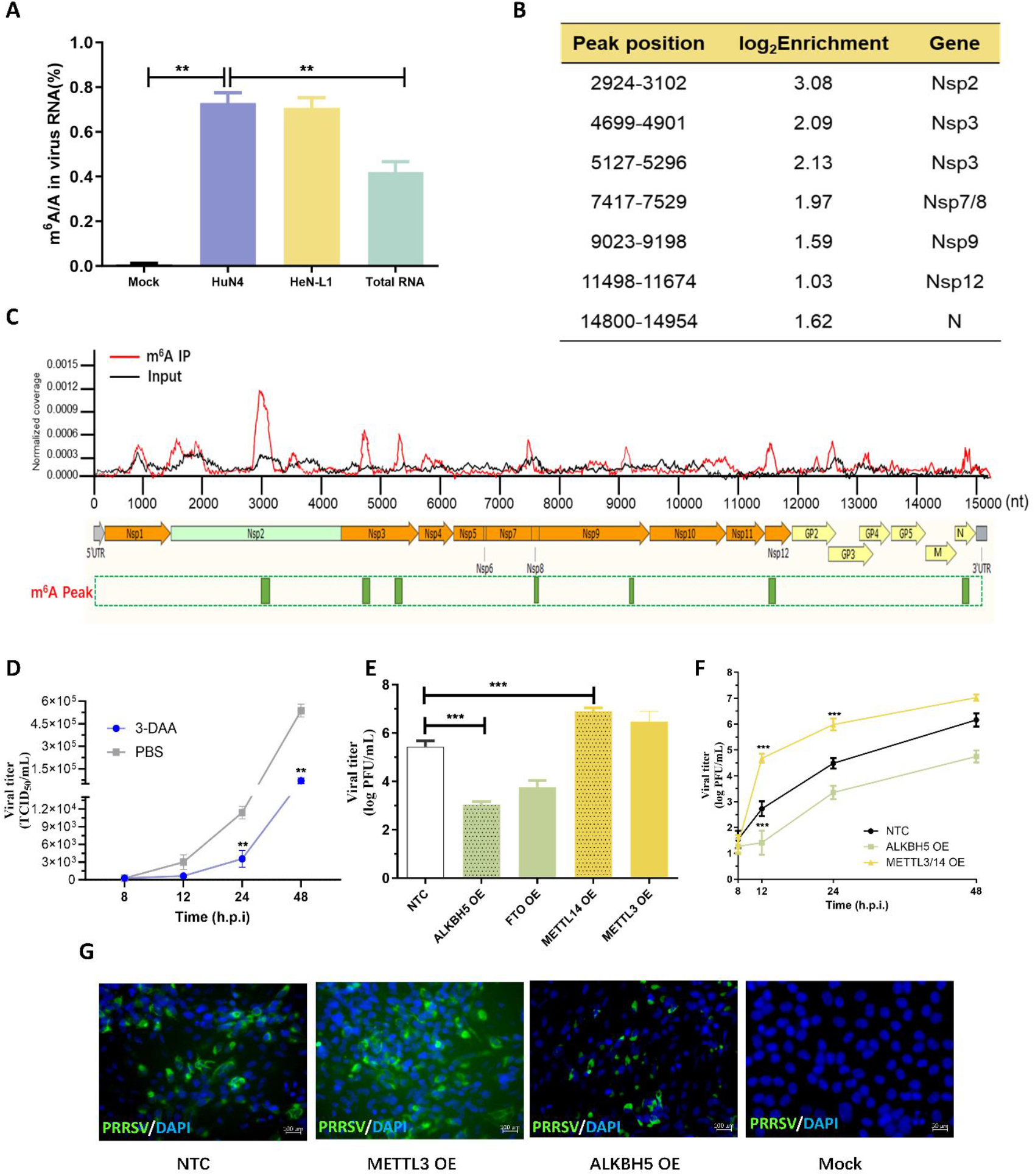
Mapping the m^6^A peaks in PRRSV genome and analyzing the correlation between m^6^A modification and PRRSV replication.

Previous studies have shown that the PRRSV genome does not encode kinases that mediating m^6^A modification. Therefore, we first interfered the expression of endogenous m^6^A related enzymes [17,18]. Marc145 cells were seeded in a six-well plate, and when the cell fusion reached 60%, 3-deazaadenosine (3-DAA) was used to inhibit the methylation of adenosine [19]. After 24 hours post transfection, the cells were then infected with PRRSV (MOI=0.1). After 8-48 hours post virus infection, the cells were subjected to repeated freeze-thaw cycles, and the virus in the supernatant was collected for TCID_50_ assay to determine the virus titer. As shown in Figure 1D, compared with PBS treated cells, 3-DAA treatment significantly decreased the virus titer from 24 to 48 hours. In order to investigate the specific effects of m^6^A enzymes on PRRSV replication, Marc145 cells were overexpressed (OE) m^6^A writers or erasers respectively. As shown in Fig 1E, overexpressed of ALKBH5 or FTO, the virus titer was significantly reduced, while in cells overexpressing (OE) METTL14 or METTL3, the virus titer was increased to approximately 10^7.0^ PFU/ml. To further validate the effect of m^6^A enzymes on the virus growth curve, Marc145 cells were seeded in a 24-well plate, and siRNA was used to interfere with the expression of ALKBH5 and METTL3/14. After transfection for 24 hours, Western blot was used to confirm the downregulation of the relevant proteins, and the cells were then infected with PRRSV (MOI=0.2). After 8, 12, 24, and 48 hours of infection, the cells were subjected to repeated freeze-thaw cycles, and the virus in the supernatant was collected for plaque assay to determine the virus titer. The cells were fixed at 24 hours, and indirect immunofluorescence assay (IFA) was used to detect the expression of PRRSV N protein. As shown in Figure 1F, compared with the backbone vector, transient overexpression of METTL3/14 enhanced virus replication, while ALKBH5 expression weakened virus replication. Similarly, IFA results in Figure 1G show that the fluorescence signal density of N protein is also affected by changes in the m6A modification level. These results preliminarily indicate that m6A modification may positively regulate PRRSV replication.

Given the profound impact of PRRSV infection on gene expression within in host cells, we used MeRIP-seq to investigate changes in m^6^A modification levels across the transcriptome of PAM cells post-PRRSV infection. As shown in Figure 2A, approximately 27,329 m^6^A peaks, among 4,677 transcripts, exhibited significant changes between HeN-L1-infected cells and uninfected cells, with m^6^A levels significantly increased in 2,258 genes and decreased in 2,419 genes. Similarly, in HuN4-infected cells, 25,641 m^6^A peaks showed significant changes, with m^6^A levels upregulated in 2,557 genes and downregulated in 2,510 genes.

**Figure 2.**
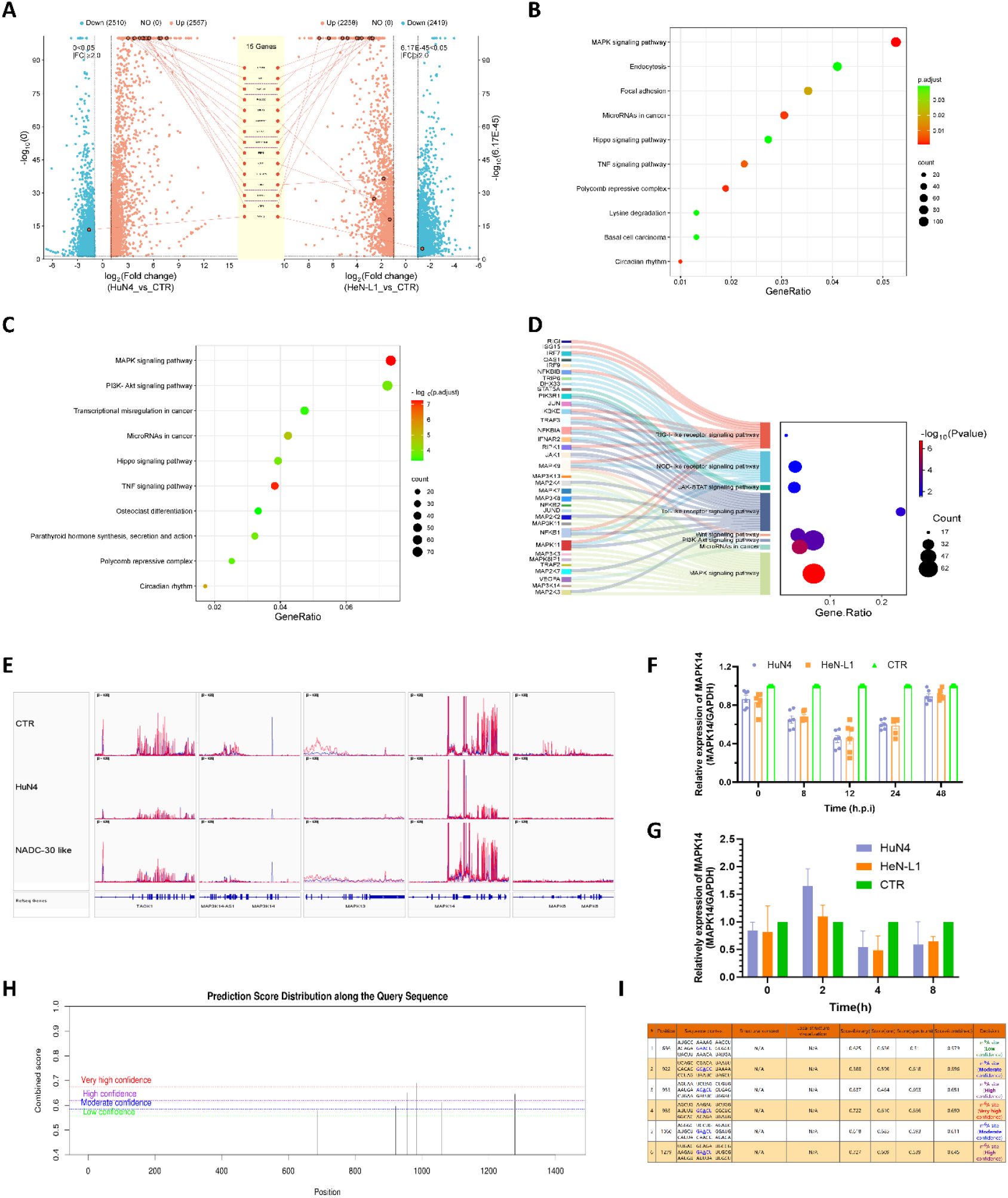
MAPK signaling pathways serving as the preliminary target of m^6^A regulating PRRSV infection.

To further explore how genes with differential m^6^A levels affect antiviral responses, we conducted the gene cluster analysis. GO and KEGG database were applied to analyze the signaling pathways associated with genes showing altered m^6^A modification levels between HeN-L1 or HuN4 infected and uninfected cells. In GO analysis (data not shown), genes with upregulated m^6^A peaks were primarily enriched in pathways like negative regulation of DNA-templated transcription, negative regulation of RNA biosynthetic process, and negative regulation of transcription by RNA polymerase II. While the results indicated m^6^A level changes in genes involved in RNA replication and cellular stress processes, further analysis is necessary to clarify the correlation between changes in m^6^A levels and specific signaling pathways. In KEGG analysis (Figure 2B and 2C), we observed a significant enrichment of MAPK signaling pathways, as well as PI3K-AKT, Hippo, TNF signaling pathways, suggesting that m^6^A modification may be crucial in modulating interactions between PRRSV and host cells. Interestingly, we screened out the genes with more than three m^6^A peaks and analyze the signaling pathways they belonging to, it came out that the pattern recognition receptors (PRRs) signaling pathways associated with antiviral innate immune response, such as RIG-I-like, NOD-like, and Toll-like receptor signaling pathways were enriched (Fig 2D). Previous studies suggest that m^6^A modification plays a key role in antiviral innate immune responses by influencing cytoplasmic PRRs recognition of viral derived nucleic acids. Our findings indicate that this mechanism may also play a significant role in PRRSV-infected PAM cells.

Among the diverse cellular signaling cascades, the p38/MAPK signaling pathway plays a particularly pivotal role due to its function as a central metabolic hub that responds to various external and internal stress stimuli. Notably, MAPK-related signaling pathways were significantly enriched in both GO and KEGG analyses in the present study, with MAPK9, MAPK11, and MAPK14 genes exhibiting multiple distinct m^6^A peaks. Mapping the m^6^A reads to the genome revealed that MAPK14 exhibited significantly different levels of m^6^A modification in both HuN4 vs. CTR and HeN-L1 vs. CTR comparisons, which indicated MAPK14 may serve as a potential critical target for m^6^A-mediated regulation of PRRSV infection (Fig 2E).

To further investigate how m^6^A modification regulates MAPK14 expression following viral infection, we first examined the dynamic changes of MAPK14 mRNA levels in PRRSV infected PAM cells. The results (Fig 2F) show that compared with the uninfected cells, both HuN4 and HeN-L1 infection in the early-phase (0-24 hours) led to a decrease in MAPK14 mRNA levels. However, from 24 to 48 hours post-infection, MAPK14 mRNA levels partially recovered, consistent with the trend of protein expression. This indicates that MAPK14 transcriptional levels are affected during the early stages of PRRSV infection. It has been known that m^6^A modification may enhance the cytoplasmic mRNA decay, therefore affecting mRNA stability. To examine whether the observed decrease in MAPK14 mRNA is associated with reduced mRNA stability due to m^6^A modification, we used actinomycin D (actD) to inhibit intracellular mRNA transcription. The results (Fig 2G) showed that fewer of MAPK14 mRNA were identified in PRRSV infected cells compared to uninfected cells, which indicating the decreased stability of MAPK14 mRNA. Therefore, we preliminarily supposed that m^6^A modification regulate MAPK14 expression by influencing its mRNA stability. To further confirm the m^6^A modification sites within MAPK14 mRNA, we used the online prediction tool SRAMP (http://www.cuilab.cn/sramp) to identify potential m^6^A modification sites. The results suggest that multiple modification sites exist in the MAPK14 gene, including three sites with high or very high confidence (Fig 2H and 2I). These findings provide a crucial foundation for further studies on how m^6^A modification regulates MAPK14 expression following PRRSV infection. In summary, our study indicates that the MAPK signaling pathway, particularly the MAPK14 gene, may be a key target through which m^6^A modification regulates the interaction between PRRSV and host cells.

In summary, the importance of m^6^A in viral infections cannot be overstated. This modification serves as a crucial player in regulating various aspects of the viral lifecycle, from RNA stability and translation to immune evasion. By studying the role of m^6^A in specific viruses, researchers can gain valuable insights into the mechanisms of viral pathogenesis and host-virus interactions. Furthermore, understanding these processes opens up new avenues for therapeutic interventions aimed at viral diseases. As our knowledge of m^6^A continues to expand, it is essential to explore its potential as a target for antiviral strategies, paving the way for innovative approaches to combat viral infections.

## Acknowledgments

This work was supported by the Central Guidance on Local Science and Technology Development Fund of Hebei Province (226Z2901G) to J.T.

## References

1. Murakami S, Jaffrey SR (2022) Hidden codes in mRNA: Control of gene expression by m(6)A. Mol Cell 82: 2236–2251.

2. Tong J, Zhang W, Chen Y, Yuan Q, Qin NN, Qu G (2022) The Emerging Role of RNA Modifications in the Regulation of Antiviral Innate Immunity. Front Microbiol 13: 845625.

3. Yu PL, Cao SJ, Wu R, Zhao Q, Yan QG (2021) Regulatory effect of m(6) A modification on different viruses. J Med Virol 93: 6100–6115.

4. Jiang X, Liu B, Nie Z, Duan L, Xiong Q, Jin Z, Yang C, Chen Y (2021) The role of m6A modification in the biological functions and diseases. Signal Transduct Target Ther 6: 74.

5. Sendinc E, Shi Y (2023) RNA m6A methylation across the transcriptome. Mol Cell 83: 428–441.

6. Chen T, Greene GH, Motley J, Mwimba M, Luo GZ, Xu G, Karapetyan S, Xiang Y, Liu C, He C, Dong X (2024) m(6)A modification plays an integral role in mRNA stability and translation during pattern-triggered immunity. Proc Natl Acad Sci U S A 121: e2411100121.

7. Dominissini D, Moshitch-Moshkovitz S, Schwartz S, Salmon-Divon M, Ungar L, Osenberg S, Cesarkas K, Jacob-Hirsch J, Amariglio N, Kupiec M, Sorek R, Rechavi G (2012) Topology of the human and mouse m6A RNA methylomes revealed by m6A-seq. Nature 485: 201–206.

8. Huang S, Wylder AC, Pan T (2024) Simultaneous nanopore profiling of mRNA m(6)A and pseudouridine reveals translation coordination. Nat Biotechnol.

9. Chen Y, Wang W, Zhang W, He M, Li Y, Qu G, Tong J (2023) Emerging roles of biological m(6)A proteins in regulating virus infection: A review. Int J Biol Macromol 253: 126934.

10. Ruedas-Torres I, Rodriguez-Gomez IM, Sanchez-Carvajal JM, Larenas-Munoz F, Pallares FJ, Carrasco L, Gomez-Laguna J (2021) The jigsaw of PRRSV virulence. Vet Microbiol 260: 109168.

11. Wensvoort G, Terpstra C, Pol JM, ter Laak EA, Bloemraad M, de Kluyver EP, Kragten C, van Buiten L, den Besten A, Wagenaar F, et al. (1991) Mystery swine disease in The Netherlands: the isolation of Lelystad virus. Vet Q 13: 121–130.

12. An TQ, Tian ZJ, Xiao Y, Li R, Peng JM, Wei TC, Zhang Y, Zhou YJ, Tong GZ (2010) Origin of highly pathogenic porcine reproductive and respiratory syndrome virus, China. Emerg Infect Dis 16: 365–367.

13. Li Y, Wang X, Bo K, Wang X, Tang B, Yang B, Jiang W, Jiang P (2007) Emergence of a highly pathogenic porcine reproductive and respiratory syndrome virus in the Mid-Eastern region of China. Vet J 174: 577–584.

14. Tong GZ, Zhou YJ, Hao XF, Tian ZJ, An TQ, Qiu HJ (2007) Highly pathogenic porcine reproductive and respiratory syndrome, China. Emerg Infect Dis 13: 1434–1436.

15. Zhou L, Wang Z, Ding Y, Ge X, Guo X, Yang H (2015) NADC30-like Strain of Porcine Reproductive and Respiratory Syndrome Virus, China. Emerg Infect Dis 21: 2256–2257.

16. Zhang H, Leng C, Ding Y, Zhai H, Li Z, Xiang L, Zhang W, Liu C, Li M, Chen J, Bai Y, Kan Y, Yao L, Peng J, Wang Q, Tang YD, An T, Cai X, Tian Z, Tong G (2019) Characterization of newly emerged NADC30-like strains of porcine reproductive and respiratory syndrome virus in China. Arch Virol 164: 401–411.

17. Liu J, Yue Y, Han D, Wang X, Fu Y, Zhang L, Jia G, Yu M, Lu Z, Deng X, Dai Q, Chen W, He C (2014) A METTL3-METTL14 complex mediates mammalian nuclear RNA N6-adenosine methylation. Nat Chem Biol 10: 93–95.

18. Roundtree IA, Evans ME, Pan T, He C (2017) Dynamic RNA Modifications in Gene Expression Regulation. Cell 169: 1187–1200.

19. Chen LJ, Liu HY, Xiao ZY, Qiu T, Zhang D, Zhang LJ, Han FY, Chen GJ, Xu XM, Zhu JH, Ding YQ, Wang SY, Ye YP, Jiao HL (2023) IGF2BP3 promotes the progression of colorectal cancer and mediates cetuximab resistance by stabilizing EGFR mRNA in an m(6)A-dependent manner. Cell Death Dis 14: 581.

